# Enhanced plasticity of programmed DNA elimination boosts adaptive potential in suboptimal environments

**DOI:** 10.1101/448316

**Authors:** Valerio Vitali, Rebecca Hagen, Francesco Catania

## Abstract

The impact of ecological changes on the development of new somatic genomes has thus far been neglected. This oversight yields an incomplete understanding of the mechanisms that underlie environmental adaptation and can be tackled leveraging the biological properties of ciliates. When *Paramecium* reproduces sexually, its polyploid somatic genome regenerates from the germline genome via a developmental process, Programmed DNA elimination (PDE), that involves the removal of thousands of ORF-interrupting germline sequences. Here, we demonstrate that exposure to sub-optimal temperatures impacts PDE efficiency, prompting the emergence of hundreds of alternative DNA splicing variants that dually embody cryptic (germline) variation and *de novo* induced (somatic) mutations. In contrast to trivial biological errors, many of these alternative DNA isoforms display a patterned genomic topography, are epigenetically controlled, inherited trans-somatically, and under purifying selection. Developmental thermoplasticity in *Paramecium* is a likely source of evolutionary innovation.

## Introduction

Developmental plasticity—the environmentally induced phenotypic variance associated with alternative developmental trajectories—has been proposed to fuel adaptive evolution by initiating phenotypic changes **(West-Eberhard 2005; Uller et al. 2018)**. Exploring the molecular mechanisms that underlie developmental plasticity can reveal a direct link between environmental changes and phenotypic differentiation, shedding light on how variation can surface from a single genotype in a stressful environment. This knowledge has important consequences for current understanding of evolutionary processes and human health **(Lea et al. 2017b; Lea et al. 2017a)**.

Previous studies in flies, plants, fungi, and vertebrates suggest that environmental changes that alter the molecular chaperone Hsp90’s buffering capacity during development can unlock cryptic genetic variation and boost phenotypic diversification **(Rutherford and Lindquist 1998; Queitsch et al. 2002; Yeyati et al. 2007; Jarosz and Lindquist 2010; Rohner et al. 2013)**. These observations substantiate an evolutionary model where cryptic developmental variation, which is revealed in response to environmental stress, might become genetically assimilated **(Waddington 1953)**. An alternative mechanism that links genetic and phenotypic variation via environmental stress has also been proposed. Recent studies in flies suggest that environmental stress, rather than exposing cryptic variation, may induce *de novo* mutations, DNA deletions and transposon insertions **(Fanti et al. 2017)**, which can result from the disruption of a class of germline-specific small RNAs known as Piwi-interacting RNAs **(Specchia et al. 2010; Gangaraju et al. 2011)**. Following stress-induced epigenetic changes, transposon activation or DNA deletions would generate somatic changes, which might ultimately become heritable via *de novo* germline mutations **(Fanti et al. 2017)**. Studying the environmental sensitivity of developmental processes across different and evolutionary distant genetic systems offers a way to test the generalizability of these cryptic and *de novo* variation-based models, possibly providing fresh insights into a molecular basis of developmental plasticity. It also provides new knowledge on the pervasiveness of phenotypic plasticity and advances current understanding of the role that environmental induction plays in adaptive evolution.

Ciliated protozoans are a biological system that enables easy manipulation of environmental conditions during development. In ciliates, nuclear development and germline-soma differentiation take place within a single cell **(Sonneborn 1977; Prescott 1994)**. Early studies in the ciliate *Paramecium* have shown that the exposure of genetically identical cells to different environmental conditions during nuclear differentiation, the so-called ‘sensitive period’, leads to heritable phenotypic variations **(Jollos 1921)**. Two of these environmentally sensitive traits have been extensively characterized. The first is the A system of complementary mating type determination, wherein the temperature to which clonal *Paramecium* cells are exposed during the sensitive period greatly influences mating type expression **(Sonneborn 1947)**. The second is the trichocyst discharge phenotype in homozygous clones of *P. tetraurelia,* where both temperature and food availability during development radically affect the phenotype expressed **(Sonneborn and M.V. 1979)**. The stable trans-generational epigenetic inheritance of phenotypes is well established in *Paramecium,* leaving open the possibility that environmentally sensitive traits may have evolutionary consequences in this ciliate. Furthermore, the mechanisms behind non-Mendelian trait inheritance in *Paramecium* are beginning to be understood at the molecular level **(Duharcourt et al. 1998; Garnier et al. 2004; Lepere et al. 2008; Duharcourt et al. 2009; Singh et al. 2014)**. Specifically, an intricate small RNA-mediated trans-nuclear crosstalk allows at least part of the genetic variability in the somatic nucleus to be inherited trans-generationally **(Coyne et al. 2012; Allen and Nowacki 2017)**. This knowledge is salient to investigations aimed at exploring the evolutionary impact that environmental changes may have on germline-soma differentiation in *Paramecium.*

Nuclear development in ciliates is coupled with a spectacular, reproducible process of selective DNA elimination from the developing somatic genome commonly known as Programmed DNA Elimination (PDE). PDE has been thoroughly characterized in *P. tetraurelia*—in addition to a broad range of eukaryotes **(Wang and Davis 2014)**, such as sea lamprey **(Smith et al. 2018b)**, finches **(Biederman et al. 2018)** or humans **(Jung et al. 2006)**. The germline genome of this ciliated protozoan comprises some 45,000 mainly unique sequences known as Internal Eliminated Sequences (IESs) **(Arnaiz et al. 2012)**. These intervening sequences are flanked by two 5’-TA-3’ dinucleotides whose disruption causes IES retention **(Mayer and Forney 1999)**, and reside both nearby and within genes, often interrupting open reading frames (ORFs). At each event of sexual reproduction, *P. tetraurelia* undergoes nuclear replacement *i.e.* the maternal somatic macronucleus is degraded and new macronuclei are produced through amplification (from 2n to ∼800n) and extensive rearrangement of the germline genome housed in mitotic copies of the zygotic nucleus **(Betermier and Duharcourt 2014)**. At this stage, IESs must be reproducibly eliminated from the germline template; their accurate and efficient splicing from genes is essential for the correct functioning of the somatic genome and the production of viable sexual offspring **(Arnaiz et al. 2012)**. Lethal developmental defects result when Piggy MAC (PGM), the domesticated transposase required for the excision of virtually all IESs and forming complexes with five additional partners **(Bischerour et al. 2018)**, is silenced **(Baudry et al. 2009; Dubois et al. 2012)**. However, viable sexual offspring are yielded when Dcl2 and Dcl3, dicer-like proteins involved in the biogenesis of small RNAs (scnRNAs) and in the excision of a subset of IESs **(Lepere et al. 2009; Sandoval et al. 2014; Hoehener et al. 2018)**, are (independently) RNAi silenced. This latter observation demonstrates that inefficient IES excision can be tolerated to some degree. Additionally, two types of erroneous DNA elimination under spontaneous conditions (in addition to variable chromosome fragmentation) were described even before IESs were comprehensively catalogued **(Duret et al. 2008; Catania et al. 2013)**: Erroneous IES excision and ‘cryptic IES recognition’. The former consists largely of inefficient excision, where IESs are excised from only a fraction of the ∼800 macronuclear copies, and to a lesser extent, of alternative or nested boundaries usage. The second type prompts the elimination of IES-like bits of somatic DNA. Therefore, although PDE is a largely reproducible developmental program for the exclusion of unwanted DNA from the somatic lineage, inaccurate DNA elimination may take place during nuclear differentiation, leading to the production of alternatively rearranged versions of the same genome.

The somatic variability that (an inefficient) PDE introduces in otherwise genetically identical *Paramecium* cells **(Caron 1992; Duret et al. 2008)** can be the basis for at least part of the phenotypic differentiation in identical clones. For example, in *P. tetraurelia* and *P. octaurelia,* an IES-like somatic region (cryptic IES) containing the promoter and the transcription start site of the mtA gene is variably spliced during development, resulting in complementary mating type determination **(Singh et al. 2014)**. A similar mechanism is found in *P. septaurelia,* a closely related species, where a cryptic IES is removed from the coding region of mtB, a putative transcription factor required for the expression of mtA, leading to mating type switch **(Singh et al. 2014)**. Thus, PDE in *Paramecium* has been repeatedly coopted for the regulation of gene expression through alternative DNA splicing **(Orias et al. 2017)**. It is conceivable that DNA-splicing-controlled phenotypes in *Paramecium* have evolved via selection of heritable alternative DNA splicing variants, consistent with previously proposed models of epigenetic evolution **(Coyne et al. 2012; Allen and Nowacki 2017)**. Although cryptic IES excision is thus far the only characterized mechanism of PDE-dependent phenotypic diversification, in principle other sources of somatic variability such as inefficient IES excision could contribute to the emergence of genetic novelties **(Catania et al. 2013; Catania and Schmitz 2015)** and adaptive phenotypic plasticity **(Noto and Mochizuki 2017; Noto and Mochizuki 2018)**.

In this study, we tested the effect that the environmental temperature has on germline-soma differentiation in *P. tetraurelia.* Our findings demonstrate, for the first time, that programmed DNA elimination in ciliates is an environmentally sensitive process. Since a large number of the IESs affected by temperature changes are epigenetically controlled and are passed down to sexual offspring, our findings also indicate that PDE is a powerful molecular ‘stonecutter’ capable of generating adaptive somatic variability.

## Results

### Temperature affects the rate of incomplete IES excision and cryptic IES recognition

After allowing isogenic lines of *P. tetraurelia* to undergo autogamy (self-fertilization) at three different temperatures, 18°C, 25°C, and 32°C, we inspected the independently rearranged somatic genomes of these lines (18°C_F1_, 25°C_F1_, and 32°C_F1_) and their progenitor (25°C_F0_) for incomplete IES excision and cryptic IES recognition. hereinafter we will refer to 18°C and 32°C as suboptimal temperatures compared to 25°C. We also arbitrarily define IES Retention Scores (IRSS) > 0.1 as non-trivial.

We detected ∼400 IES loci with IRS > 0.1 (hereinafter, somatic IESs) in the progenitor line 25°C_F0_ **(Figure 1A)**. This count is comparable to that estimated for the descendant 25°C_F1_ and remains similar between the two 25°C samples, but it changes with the irs threshold applied—note that the IRS is unaffected by between-sample variation in read coverage. In contrast, the count of somatic iess is considerably higher in the macronuclear genomes that developed at suboptimal temperatures. we detected up to ∼800 somatic iess in macronuclear genomes rearranged at 18°C and 32°C, roughly a two-fold increase compared to the optimal temperature. the vast majority of these somatic iess have scores in the range of 0.1-0.3 **(Figure 1a)**, with a potential impact on gene expression.

**Figure 1.**
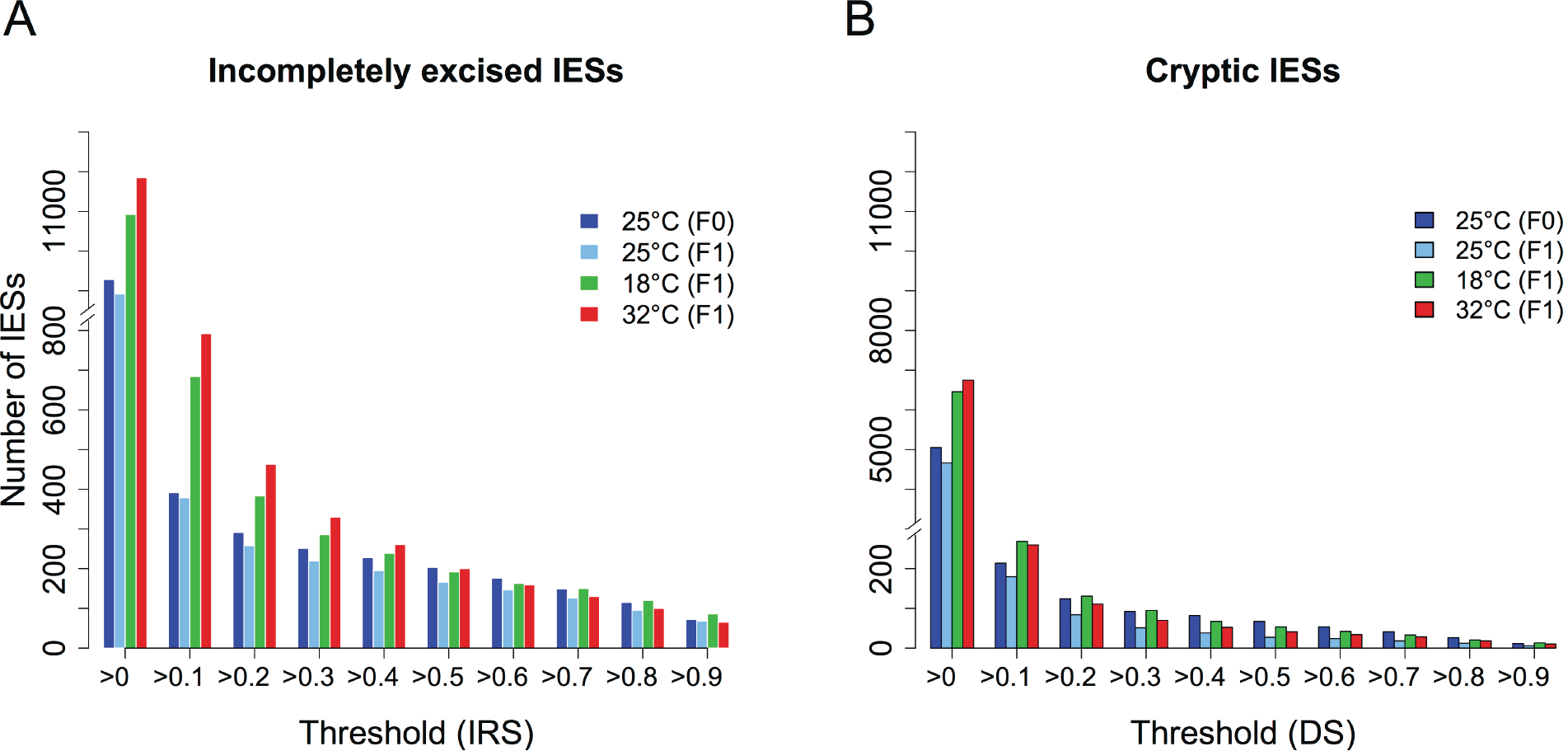
The rate of incomplete IES excision and somatic DNA deletion is temperature dependent. (A) The number of detected incompletely excised IESs is higher at 18°C and 32°C compared to 25°C. For non-trivial IES retentions (IRS > 0.1), the mean count bottoms at ∼400 IESs at 25°C, whereas it rises up to ∼750-800 IESs at suboptimal growth temperatures (18°C and 32°C). (B) TA-bound somatic deletions (Cryptic IESs) are also more frequent when PDE occurs at suboptimal growth temperatures. IESs, Internal Eliminated Sequences; PDE, Programmed DNA Elimination; IRS, IES Retention Score; DS, Deletion Score.

Suboptimal temperatures affect also the rate of cryptic IES recognition (TA-bound somatic deletions) **(Figure 1B)**. We detect up to a ∼2-fold increase when comparing the number of partially excised cryptic IESs unique to 18°C and 32°C with those unique to the 25°C samples **(Figure S1)**. Many TA-bound somatic deletions lead to partial or even complete gene ablation and occasionally span multiple genes at once (a catalogue is presented in Table S1). Among the 18 TA-bound somatic deletions consistently retrieved across all temperatures (maternally inherited), we find the 195 bp-DNA segment containing the promoter and transcription start site of mtA, a DNA-splicing regulated gene **(Singh et al. 2014; Orias et al. 2017)**. We also report a set of IES-like somatic regions that are variably spliced at different temperatures, which might represent temperature-sensitive cryptic IESs. The surge in cryptic IES deletions with temperature is IESs pronounced compared to that in the annotated IESs and mostly limited to deletion scores (DS) that tend to be smaller than 0.1 **(Figure 1B)**. As a consequence, we decided to focus the presentation of our results on true IESs.

### Suboptimal environmental temperatures decrease PDE efficiency

We investigated how extensively temperature changes affect the magnitude of IES retention in the polyploid somatic genome. The full set of PGM-IESs with their retention scores (IRS), and the genome-wide testing of the F0-to-F1 IRS transitions for all samples are reported in **Table S2.**

We detected a marked reduction in PDE efficiency at 18°C and 32°C relative to 25°C **(Figure 2)**. The number of IESs with a greatly intensified retention in the F1 somatic nuclei (IRS_F1_ ≫ IRSF_0_, Binomial test, *P*_adj_ < 0.05) rises from 43 at 25°C, to 183 and 271 at 18°C and 32°C, respectively. Further, most of the significantly retained IESs detected at 18°C and 32°C are unique to sub-optimal temperatures—with a treatment-control ratio of ∼12-fold (151:13) and ∼17-fold (225:13) for 18°C and 32°C, respectively (Figure S2A). Yet the number of IESs excised with significantly *increased* efficiency in the F1 generation is comparable across temperatures **(Figure 2)**, with around 50% overlap between each of the experimental lines and the control **(Figure S2B)**. The count of significant transitions for the three temperatures tested are summarized in **Figure 2D**. The count ratio of upward (reduced excision efficiency) to downward (increased excision efficiency) IRS transitions after sexual reproduction approaches 1 (43:57) for the control temperature of 25°C, whereas it rises up to ∼3.5 (183:54) and ∼5.5 (271:50) at 18°C and 32°C, respectively. These observations demonstrate that rather than producing stochastic effects, in our study suboptimal temperatures substantially reduce PDE efficiency, with the rate of erroneous excision rising well above biological noise.

**Figure 2.**
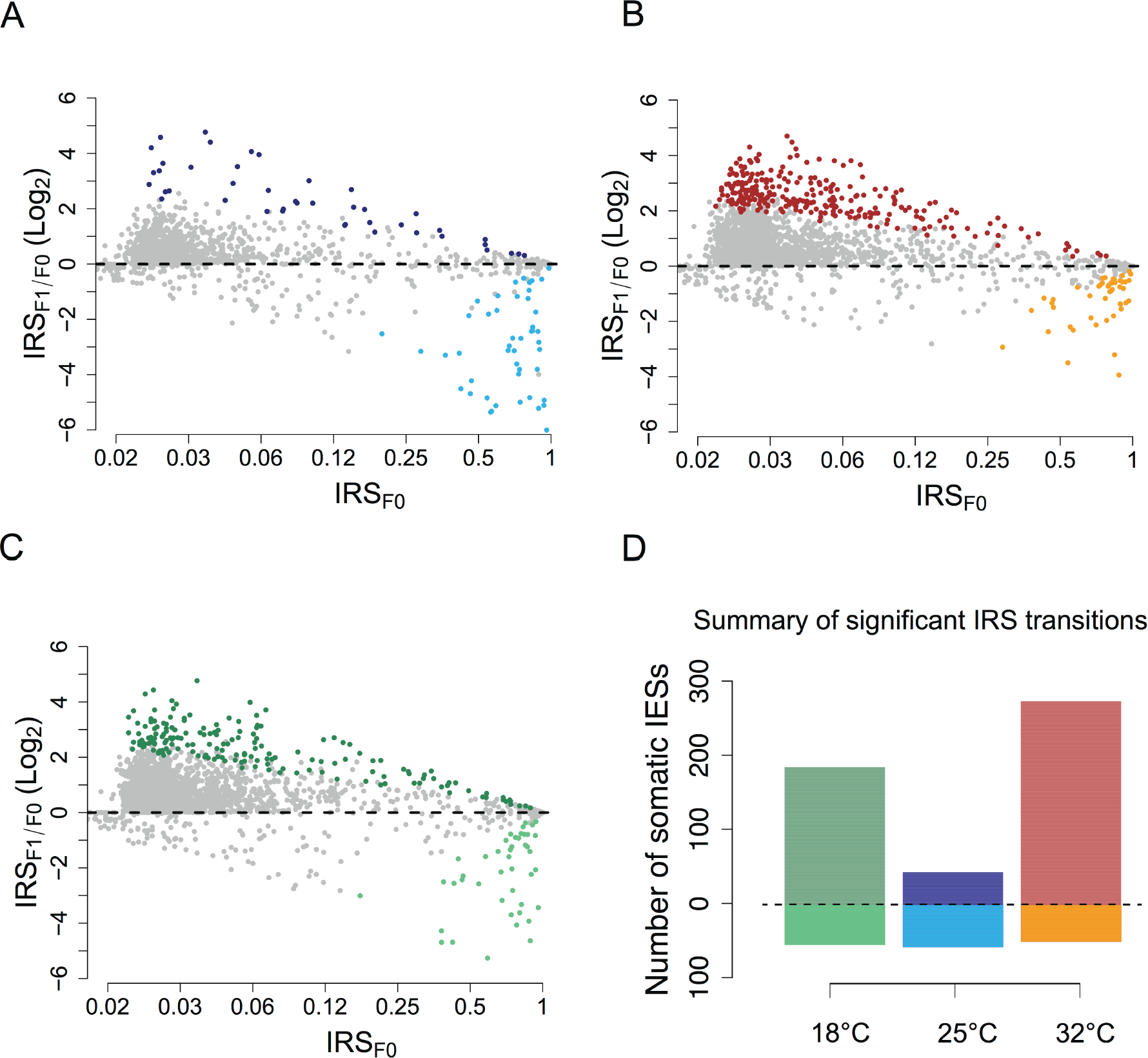
PDE inefficiency shows a characteristic U-shaped relationship with temperature. (A-C) Bland-Altman plots displaying the Log2 fold change of IRSs from F0 to F1 for 25°C, 32°C and 18°C, respectively. Statistically significant IRS transitions (Binomial test, P_adj_ < 0.05) are shown as colored filled-circles: 25°C, dark and light blue circles; 32°C, red and orange circles; 18°C, dark and light green circles (dark color: IRS_F1_ > IRS_F0_; light color: IRS_F1_ < IRS_F0_). Position on x-axis reflects the log-transformed initial (F0) state of the IRSs (x-labels are IRSs before log transformation). (D) Counts of statistically significant IRS transitions (P_adj_ < 0.05) after nuclear differentiation at 18°C (green), 25°C (blue) and 32°C (red). The number of somatic IESs experiencing an upward or downward IRS transition is shown above and below the horizontal dashed line, respectively.

### Incomplete IES excision is trans-generationally inherited

Until now, we have mainly focused on the performance of PDE at different environmental temperatures. We now turn to the potential biological significance of the PDE-mediated molecular variation. To begin, we asked whether the somatic IESs detected in the F1 genomes were all generated anew or if they were partly obtained through trans-somatic inheritance.

To gain insight into this question, we assessed how many IESs are retained simultaneously in the four independently rearranged F0 and F1 somatic nuclei. If incomplete IES excisions are truly stochastic events as commonly regarded, then the observed number of 4-way-shared IESs should not exceed what would be expected by chance. Leveraging 100 simulated datasets based on random draws from the PGM-set (*i.e.,* the set of IESs retained after Piggy MAC silencing), we estimated that the expected maximum number of IESs shared among four genomes is ∼107. We observed 934 4-way-shared IESs, a striking ∼ninefold increase **(Figure 3A)**.

**Figure 3.**
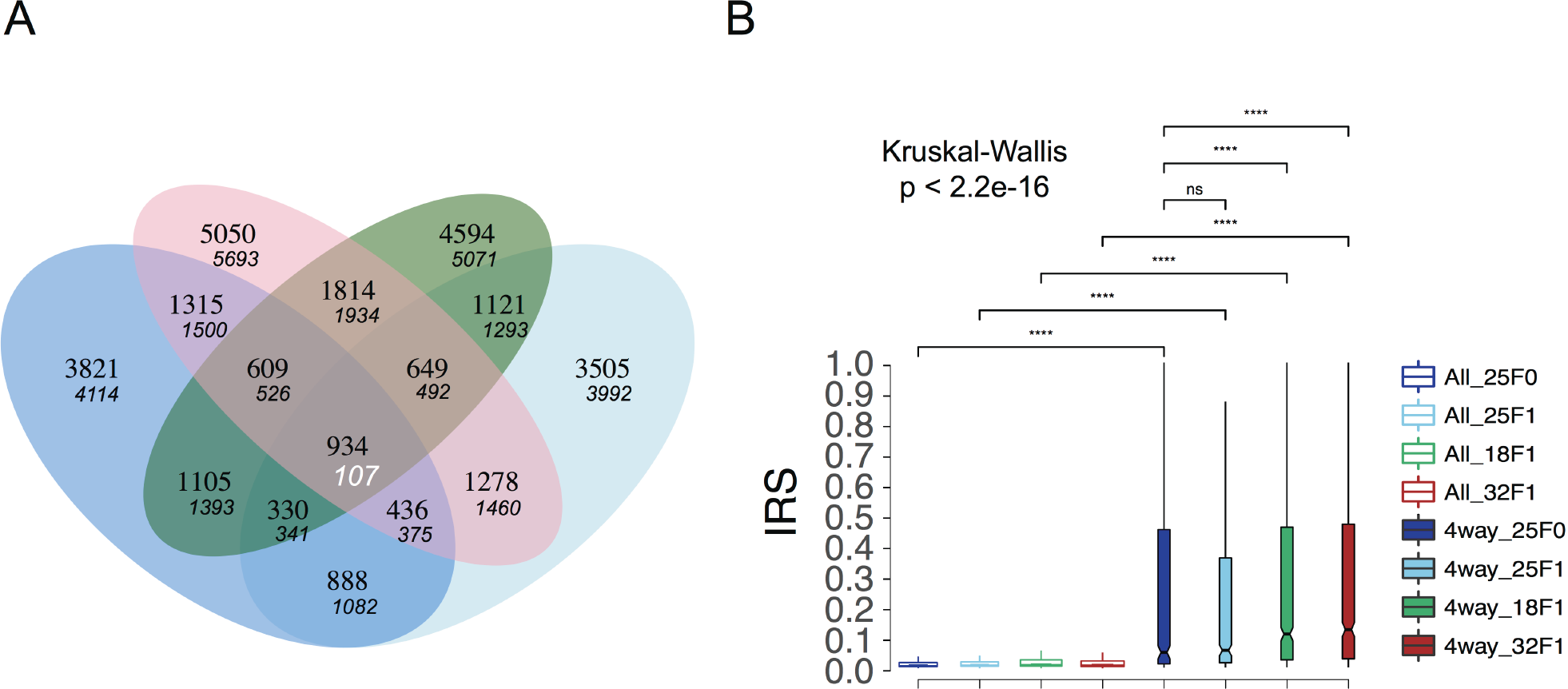
Incompletely excised IESs are inherited trans-generationally and show elevated retention scores. (A) Venn diagram depicting sets of incompletely excised IESs shared between three F1 somatic genomes and their parental F0 genome. A large excess of incompletely excised IESs is shared across all four genomes, indicating that most of the IESs in the 4-way set were inherited transsomatically from F0 to F1. The maximum number of shared IESs, as determined over 100 random draws, is shown under each observed overlap value. Blue ellipse, 25°C (F0), green ellipse, 18°C (F1), light blue ellipse, 25°C (F1), red ellipse, 32°C (F1). (B) Box plot showing the IRS distributions (IRS > 0) for the full-set and the 4-way set (4w) of incompletely excised IESs at all investigated temperatures. Pairwise comparisons between groups were performed using a Wilcoxon Rank Sum Test with correction for multiple testing (BH, Benjamini-Hochberg). Statistical significance is indicated for each comparison (****; *P* < 0.001, ns, non-significant). Outliers are omitted for clarity.

In principle, somatic IESs shared between independent macronuclei could reflect weak cis-acting IES recognition/excision signals. Weak splicing signals might explain the significantly elevated median IRS (up to complete retention, IRS ∼1) of the 4-way-shared IESs relative to the full-set of incompletely excised IESs illustrated in **Figure 3B**. Consistent with the weak-signal hypothesis is the study of the C_in_-score— a predictor of splicing signal quality **(Ferro et al. 2015)**—which reveals statistically smaller C_in_-score estimates for 4-way-shared IESs relative to the PGM-set (IESs_4-way_ vs. IESs_PGM_, 0.54 vs 0.62, Mann-Whitney U, *P* < 0.001). This difference holds true when we control for size class and genomic location. Nonetheless, weak splicing signals alone cannot explain why the same IESs exhibit significantly elevated median IRS at 32°C and 18°C relative to 25°C **(Figure 3B)**.

Another (not mutually exclusive) explanation for the excess of somatic IESs common to subsequent sexual generations is that these somatic IESs might reflect episodes of possibly ongoing trans-generational epigenetic inheritance. Under these circumstances, we expect that many of the discussed somatic IESs (including the 4-way-shared IESs) be epigenetically regulated. We tested this hypothesis taking advantage of published knock down (KD) studies of PDE-associated epigenetic components in *P. tetraurelia* **(Sandoval et al. 2014)**. Three Dicer-like endonucleases and two classes of developmentally specific small RNAs were shown to be necessary for the accurate excision of a few thousands of *P. tetraurelia* IESs: Dcl2 and Dcl3 are required for the biogenesis of scnRNAs in the germline nucleus **(Lepere et al. 2009)**, whereas Dcl5 is responsible for the production of iesRNAs in the developing somatic nucleus **(Sandoval et al. 2014)**. First we examined the set(s) of somatic IESs that are significantly retained, without being necessarily shared between independent macronuclei. We found that the excision of roughly half of these IESs is dependent on Dcl2/3 and or Dcl5, *i.e.,* their retention score increases significantly upon Dcl2/3 and Dcl5 KD **(Figure 4)**.

**Figure 4.**
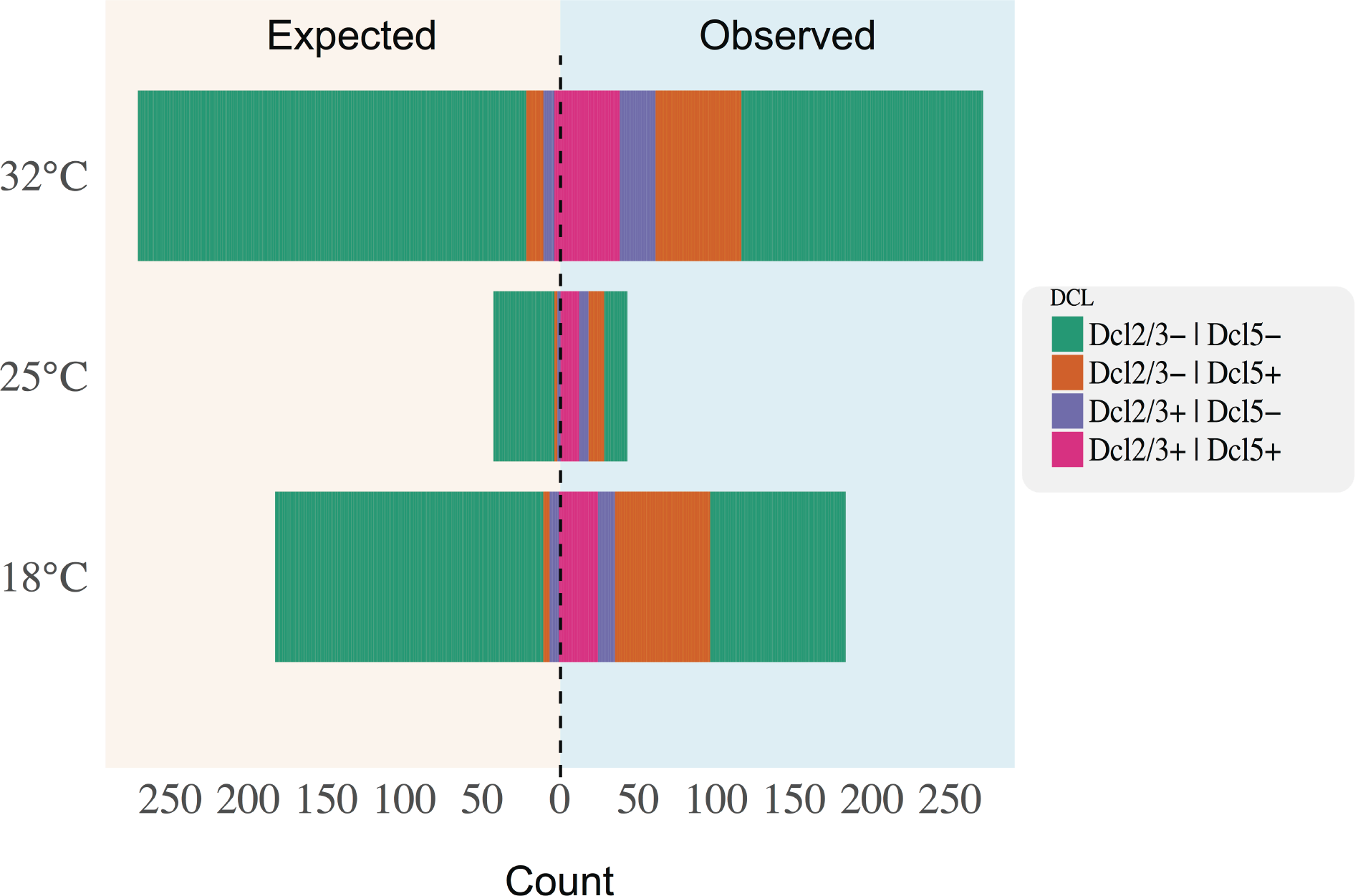
A large excess of incompletely spliced IESs is epigenetically regulated. Back to back stacked bar chart showing the number of significantly retained IESs after nuclear differentiation at all investigated temperatures. For each temperature, IES counts are broken down into Dcl2/3-controlled IESs (Dcl2/3^+^ | Dcl5^-^, Purple), Dcl5-controlled IESs (Dcl2/3^-^ | Dcl5+, Orange), Dcl2/3-Dcl5-co-controlled IESs (Dcl2/3^+^ | Dcl5^+^, Fuchsia) and Dcl-independent IESs (Dcl2/3^-^ | Dcl5^-^, green). Expected proportion of Dcl-dependent IESs for random samples of the same size (left side) are shown back to back with the observed data (right side).

We then examined how extensively the 4-way-shared IESs are under the control of Dcl2/3-5 (data taken from ParameciumDB). We found that 166 (out of the 934) IESs are indeed Dcl2/3-controlled IESs and 248 are Dcl5-controlled. This establishes that at least 35% of the 4-way-shared somatic IESs are under epigenetic control, a ∼3.6-fold enrichment compared to random expectation (∼10%), and thus likely to be epigenetically inherited.

In fact, the number of 4-way-shared incompletely excised IESs that are epigenetically controlled/inherited might be even larger. Under the scnRNA model of trans-nuclear comparison **(Duharcourt et al. 2009)**, IESs that are severely retained in the maternal macronucleus will be less affected by the scnRNA depletion ensuing Dcl2/3 KD compared to IESs that are absent from the maternal macronucleus. Conventional approaches may fail to classify these highly retained IESs as Dcl2/3-dependent because the shifts between pre-and post-KD levels of IES retention might be negligible. Leveraging the IRSs obtained by previous Dcl2/3 KD experiments **(Lhuillier-Akakpo et al. 2014; Sandoval et al. 2014)** we identified 236 IESs that despite having particularly elevated IRSs in these KD experiments (IRS > 0.3) were not recorded as influenced by the scnRNA machinery. Around 68% of these IESs (n=160) are found in our 4-way-shared set. Thus, our approach based on perturbation of environmental conditions permits the identification of trans-somatically inherited IESs without requiring any previous knowledge of the epigenetic factors involved in their excision, adding hundreds of candidate epigenetically-controlled IESs to the existing set.

Collectively, our observations suggest that a considerable number of somatic IESs in *P. tetraurelia* are trans-somatically inherited, and that the retention levels of these inherited IESs are enhanced in the sexual progeny upon exposure to suboptimal temperatures.

### Purifying selection shapes IES retention profiles

To deepen our understanding of the biological significance of somatic IESs, we performed a systematic analysis of their genomic distribution. The expectation is that non-trivially retained IESs are more prevalent in i) intergenic regions and ii) weakly expressed genes or genes that are not critical for development or cell viability.

Indeed somatic IESs (IRS > 0.1) preferentially occupy intergenic regions **(Figure 5A)**. Furthermore, they are more likely to occur in genes that in the *P. tetraurelia* strain 51 are weakly expressed in the vegetative stage **(Figure 5B)**. Interestingly, the ratio of intergenic to intragenic somatic IESs is IRS-dependent. The proportion of intergenic somatic IESs increases abruptly as the IRS crosses 0.1 **(Figure 5A)** and plateaus at ∼70% beyond an IRS threshold of ∼0.25. This is true at all the investigated temperatures.

**Figure 5.**
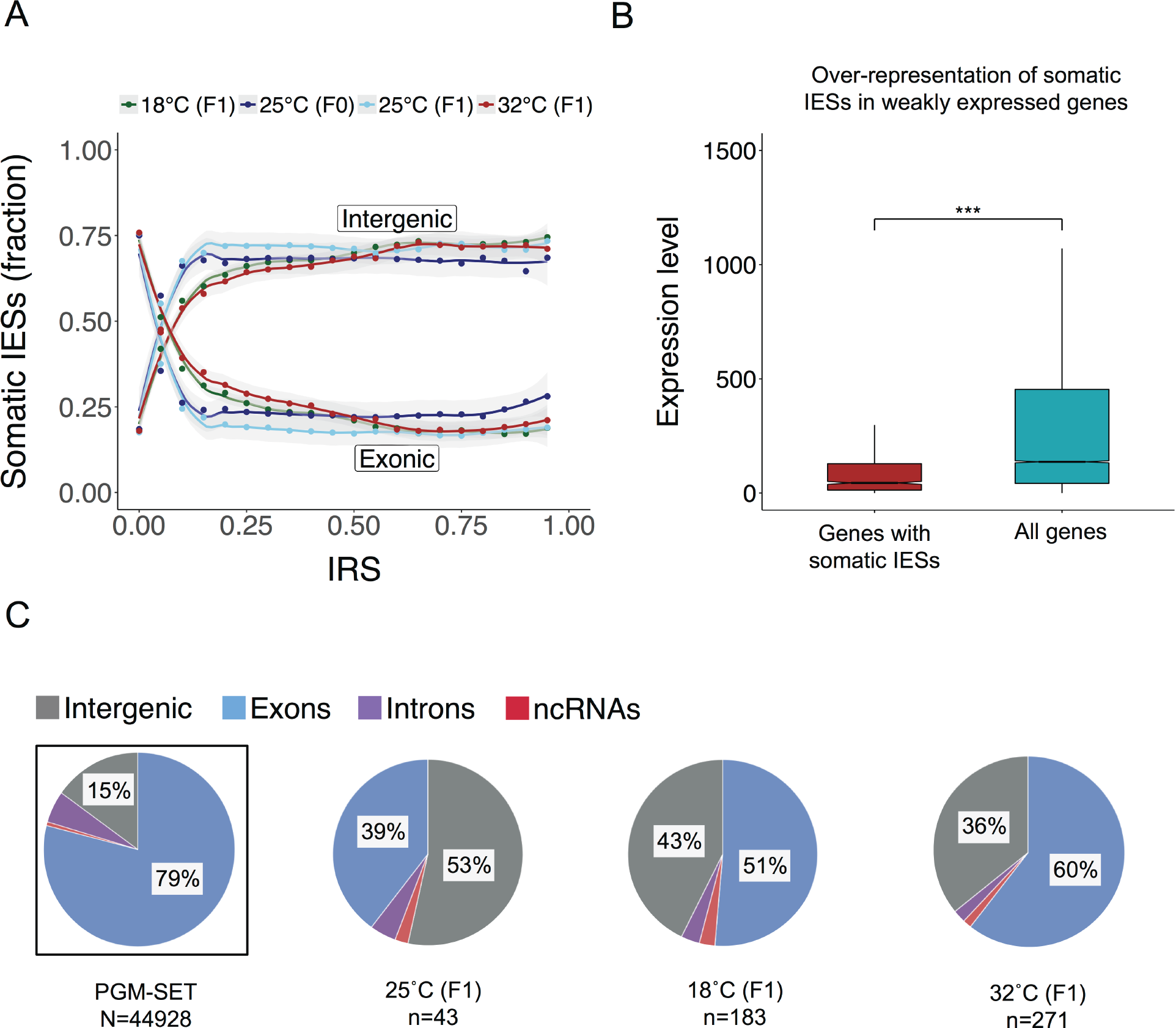
Non-trivially retained IESs are under purifying selection. (A) IRS-dependent changes of nuclear prevalence for incompletely excised intergenic and exonic IESs. (B) Expression levels distributions of genes affected by IES retention (IRS > 0.1, red box) and the full set of *P. tetraurelia* macronuclear coding genes (cyan box). Pairwise comparison was performed with a Mann-Whitney *U* test. Statistical significance is indicated (***; *P* < 0.01). (C) Genomic distribution of significantly retained IESs at all investigated temperatures. Distribution of PGM-controlled IESs is shown for reference (leftmost pie chart, boxed). Sample size is indicated in brackets. Percentages below 5% are not indicated.

When we compared the subset of significantly retained IESs (see *Suboptimal environmental temperatures decrease PDE efficiency*) with the PGM-set, we once again detected an excess of intergenic loci **(Figure 5C)**. Nevertheless, a cross-sample comparison reveals a substantial enrichment of somatic IESs within exons at 18°C and 32°C relative to 25°C **(Figure 5C)**. In particular, 83 and 158 genes show significant levels of IES retention at 18°C and 32°C, respectively, compared to only 17 genes at 25°C **(Figure S3)**. Intriguingly, the deviation from the reference distribution is for the most part determined by the 4-way-shared set of IESs. After the exclusion of these largely epigenetically-controlled IESs, the genomic distribution of the remaining set of retained IESs conforms to the expected reference distribution (data not shown).

In sum, although IESs in intergenic regions are generally more likely to be incompletely excised, suboptimal environmental temperatures appear to mostly perturb the excision of exonic, epigenetically regulated, and presumably trans-somatically inherited IESs.

### IES retention in exons disrupts ORFs largely, but not exclusively

Somatic IESs within genes could give rise to functional alternative isoforms. But how likely is this event to occur? Within coding sequences (CDS), IES retention can induce Premature Termination Codons (PTCs), either through within-IES PTC (IES-PTC) or more commonly via PTC induction downstream to the inserted element (FrameShift-PTC or FS-PTC), likely contributing to transcript degradation via Nonsense-mediated mRNA Decay (NMD) **(Brogna and Wen 2009)**. In addition, IES insertion near the 3’ end of the CDS can lead to the ablation of the true stop codon (Tail-FS IESs), presumably resulting in mRNA degradation via non-stop mediated RNA decay **(Frischmeyer et al. 2002; Vasudevan et al. 2002; Klauer and van Hoof 2012)**. Nevertheless, productive alternative DNA splicing variants could, at least in theory, be achieved via retention of non-PTC containing 3n-IESs (length multiple of 3).

To assess the impact of incomplete IES excision on genes, we calculated the occurrence of PTC-inducing and non-PTC inducing IESs in the PGM-set (control set) and compared it with their counterpart in the experimental 32°C set (which contains the largest number of significantly retained IESs). We find that ∼20% of IES retentions may in theory produce protein diversification, although only ∼5% of the IESs retained in the experimental set represent cases of theoretically productive alternative DNA splicing (3n-IESs; **Figure 6)**. Of note, Tail-FS IESs outnumber 3n-IESs in the 31 genes of the experimental set with *bona fide* CDS extension (15.8% vs. 5.4%), whereas the reverse pattern is found for the control set (3.3% vs. 20.6%). We infer that the vast majority of incomplete IES excisions would likely impact protein availability—a condition that by silencing some genes rather than others might advantageously facilitate adaptation to a new environment—although cases of potentially productive, IES-driven protein diversification may occur at each event of sexual reproduction.

**Figure 6.**
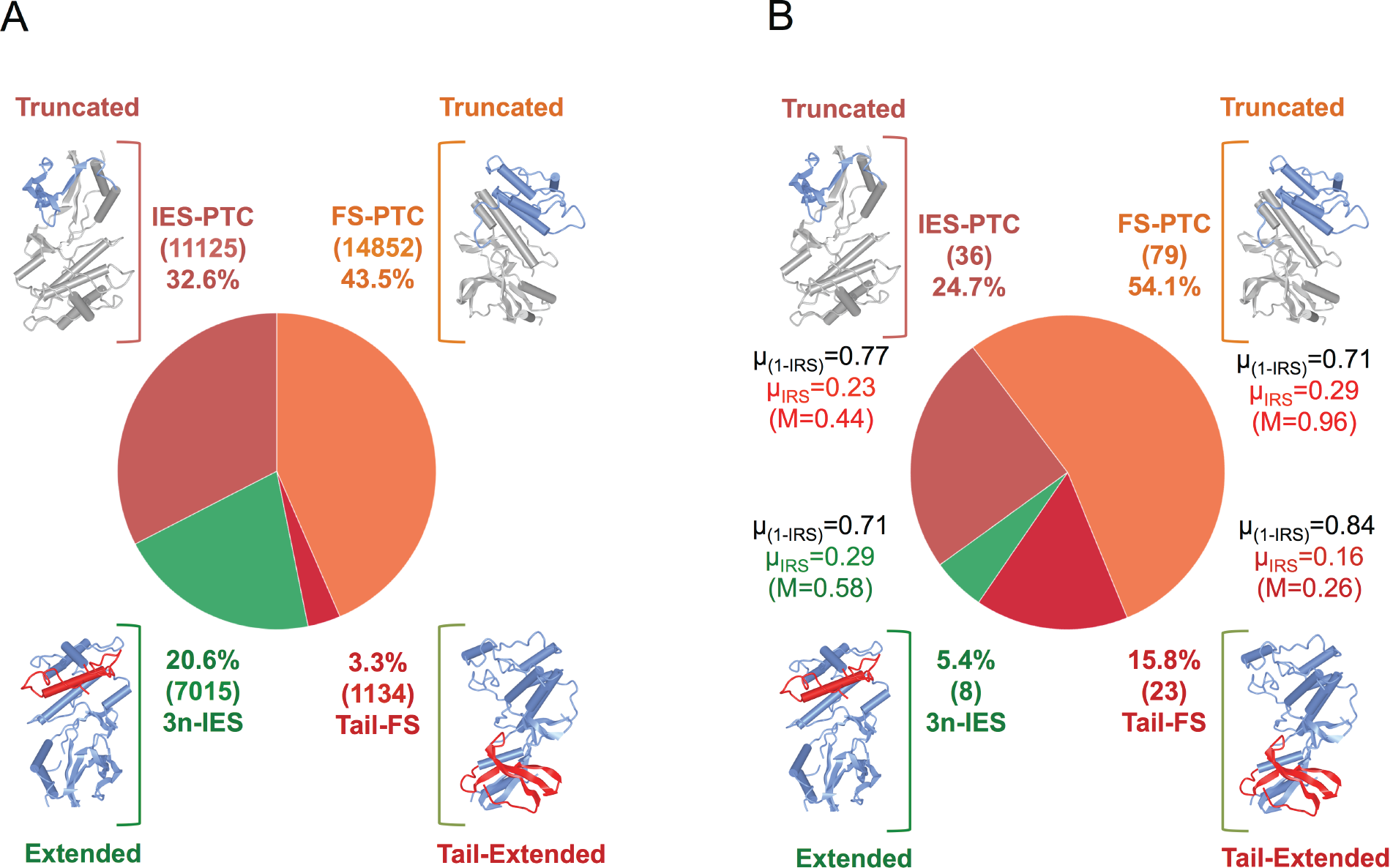
IES insertion within CDS potentially diversifies protein sequences. (A) Theoretical diversification potential of PGM-IESs. 3n-IES (green): fraction of productive, non-PTC inducing IESs. Tail-FS (crimson red): fraction of IESs that do not introduce a PTC but lead to the ablation of the true stop codon. FS-PTC (orange): fraction of IESs that disrupt the open reading frame by introducing a downstream PTC. IES-PTC: Fraction of IESs that introduces a PTC within the insertion. (B) Classification of the 146 within-CDS IES insertions observed at 32°C based on the predicted transcriptional outcome. Mean excision score (1-μIRS), Mean IRS (μIRS), Maximum IRS (M) are indicated next to each category. The number of IESs is indicated within brackets. The predicted effect on the protein products is depicted schematically next to each class. IES, Internal Eliminated Sequence; CDS, Coding Sequence; PTC, Premature Termination Codon; PGM, Piggy Mac transposase; FS, Frame-Shift.

### Non-random distribution of IESs with respect to protein families and molecular function

Finally, we explored possible biases in the molecular function of the genes with significant IES retention in coding sequences. The functional categorization of these genes is given in Figure S3. The full set of genes, their expression values (as in strain 51; taken from **(Arnaiz et al. 2017)**), and annotations, along with parameters related to the retained IESs are presented in **Table S2**.

A GO term enrichment analysis reveals a single molecular function term, *protein binding,* enriched at both 18°C and 32°C (Fisher exact test, *P* < 0.0001), but not at 25°C (P = 0.05). Interestingly, two sets of proteins, the Tetratrico Peptide Repeat Region (TPR, IPR019734) and Growth Factor Receptor cysteine-rich domain (GFR, IPR009030) containing proteins contribute largely to this functional enrichment. More specifically, TPR and GFR-genes together account for ∼21 % and ∼24% of the genes affected at 32°C and 18°C, respectively. This indicates that in *P. tetraurelia* there may be genes that are more susceptible to incomplete IES excision than others.

We hypothesized that there are genes in *P. tetraurelia* that are particularly IES-dense and thus more likely to be among the genes hit by IES retention. In testing this hypothesis, we found that genes significantly affected by IES retention at 18°C and 32°C do have significantly greater than average number of IESs **(Figure 7A)** and IES density **(Figure 7B)**. Furthermore, TPR motif-containing proteins (IPR019734) and GFR cysteine-rich domain-containing proteins (IPR009030) exhibit significantly elevated numbers of IESs *per* gene, being among the most IES-rich genes in the *P. tetraurelia* genome **(Figure 7C)**. While both the GFR and TPR protein families are extremely IES-rich, only the latter is also ultra-IES dense **(Figure 7D)**: with IES densities up to 10 IES/kb, the TPR-motif family of proteins alone accounts for almost 3% (1,200 IESs) of the 44,928 PGM-set of IESs (an example of TPR-motif gene is in **Figure 7E)**.

**Figure 7.**
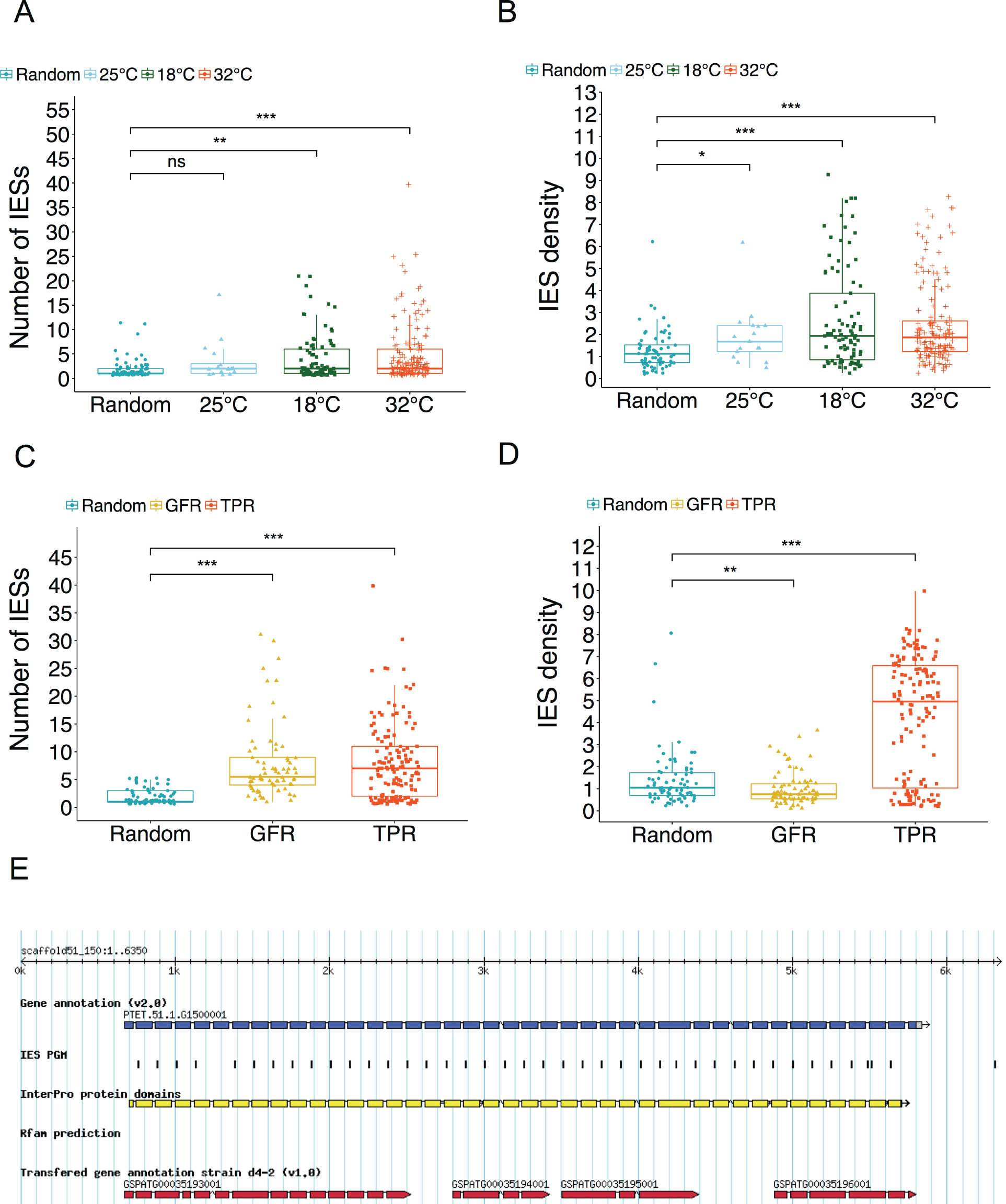
TPR-motif and GFR-Cys-rich domain containing proteins are lES-rich gene families susceptible to inefficient IES excision in response to temperature changes during autogamy. (A) Genes affected by IES retention at 18°C and 32°C have significantly greater than average number of IESs. (B) Across all the investigated temperatures, genes hit by significant IES retention are IES-dense (Kruskal-Wallis, *P* < 0.0001). (C) TPR and GFR proteins exhibit extraordinary per gene IES counts (Kruskal-Wallis, *P <* 0.0001). (D) TPR but not GFR proteins are characterized by elevated IES density (Kruskal-Wallis, *P* < 0.0001). (E) A TPR-motif gene (PTET.51.1.G1500001) showing a characteristic pattern of IES distribution. This gene is hit by multiple IES retention at 32°C. IESs are positioned almost invariably at the 5’ end of the TPR coding exons. According to the v2.0 annotation of macronuclear gene models, PTET.51.1.G1500001 is the gene with the greatest number of IESs to be found in the *P. tetraurelia* genome. TPR, Tetratrico Peptide Repeat region motif (IPR019734). GFR-Cys-rich, Growth Factor Receptor cysteine-rich domain (IPR009030). IES, Internal Eliminated Sequence. IES density, number of IESs per kb.

The pronounced representation of GFR-and TPR-containing proteins in our dataset might be merely expected by chance. To address this question, we partitioned *P. tetraurelia* genes into three groups on the basis of their InterPro domain annotations, TPR, GFR and Protein kinase-like domain (PKD) genes, with the latter group serving as a control in the enrichment analysis. We find that the number of GFR-genes (as well as PDK-genes) does not differ significantly from the expected values (see Materials and Methods), neither at 32°C nor at 18°C (binomial test, *P* >0.05). Conversely, TPR-genes are highly overrepresented at both sub-optimal temperatures (binomial test, *P* <0.0001). Thus, TPR-genes in *P. tetraurelia* appear to be highly susceptible to IES retention at sub-optimal temperatures.

Guided by this non-random distribution of IESs in relation to protein families, we next asked whether *P. tetraurelia* IESs are generally non-randomly distributed with respect to molecular functions or biological processes. To address this question, we performed enrichment analysis on the subset of *P. tetraurelia* IES-containing genes using the full set of *P. tetraurelia* macronuclear genes as reference. Remarkably, 10 molecular function and 8 biological process terms are enriched in the set of IES-containing genes **(Table 1)**. This suggests that *P. tetraurelia* genes involved in specific cellular functions such as ion transport, signal transduction and microtubule-based movement, among others, are relatively more likely to contain IESs. Conversely, we find a deficit of IESs in e.g. translation-associated factors.

**Table 1.**
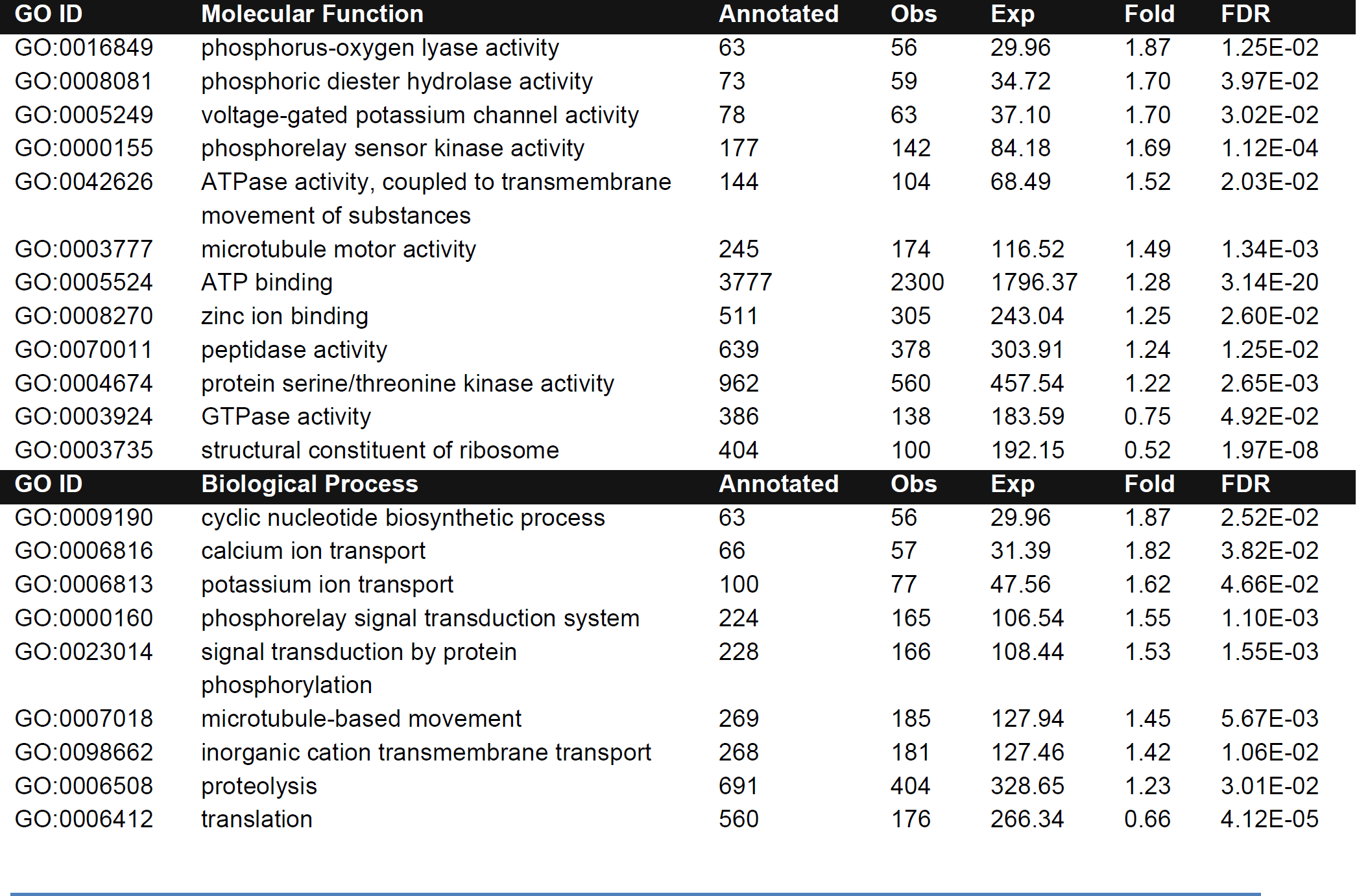
IES-containing genes are enriched in specific molecular functions and biological processes. GO ID, GO term IDs. Annotated, number of genes mapped to the corresponding GO term in the genome. Obs, Observed number of genes; Exp, Expected number of genes. Fold, Fold Enrichment of the functional category. FDR (False Discovery Rate), Fisher’s exact test with Benjamini-Hochberg correction for multiple testing, *P_adj_* < 0.05. The results shown in the table were obtained using the *Panther gene list analysis* tool. Similar results were obtained performing the functional enrichment analysis with the *topGO* package (not shown). The outcome of the GO-Term analysis was highly consistent between gene annotation versions (macronuclear gene models v1 and v2).

These findings demonstrate that IESs are not randomly distributed in relation to protein families, molecular functions, and biological processes. The functional enrichment of IES-containing genes could be explained in terms of i) a spatially patterned genomic invasion by transposable IES progenitors due to heterogeneous levels of purifying selection that antagonizes gene interruption and/or ii) differential expansion of gene families in the *P. tetraurelia’s* genome following the invasion. An alternative, tempting explanation is that IESs’ patterned genomic topography has been shaped by natural selection over evolutionary time to help regulate organismal responses to environmental changes.

## Discussion

In this work we asked: How extensively might environmental changes affect germline-soma differentiation? Answering this question contributes to our view of how the crosstalk between genes and environment affects organismal constitution and evolution. One possibility is that environmental changes may have generally little impact on this developmental process. Alternatively, environmentally induced perturbations might introduce considerable molecular variation into the developing somatic genome. Our results provide support for the second perspective. They also suggest that environmentally induced somatic variation in *Paramecium* might be evolutionarily relevant.

We show that the rate of spontaneous IES retention and cryptic IES recognition increase at sub-optimal temperatures **(Figure 1)**, with the mean error rate of IES excision with IRS > 0.1 rising from ∼200 at 25°C to ∼600 erroneously excised IESs per sexual generation. In addition, the inefficacy of PDE during *P. tetraurelia’s* autogamy has a characteristic U-shaped relationship with temperature—the inefficiency of IES excision peaks at both low (18°C) and high temperatures (32°C) whereas PDE experiences an optimal performance at 25°C **(Figure 2)**. IES excision is therefore greatly sensitive to changes in the environmental temperature, a finding that may be surprising given that *Paramecium* is continuously exposed to quotidian and seasonal temperature fluctuations in naturally occurring conditions, thriving and undergoing sexual reproduction along latitudinal temperature gradients **(Krenek et al. 2011; Krenek et al. 2012)**. In as little as one sexual generation, the thermo-plasticity of IES excision translates into the introduction of hundreds of new alternative DNA splicing variants and in an elevated nuclear prevalence of standing somatic variation.

What is the cause of this increased somatic genetic variation? Intracellular processes coupled with DNA repair such as meiotic recombination have been recently shown to be thermo-plastic, with a similar U-shaped temperature-performance response **(Lloyd et al. 2018)**. Much like meiotic recombination, both biophysical and physiological alterations in response to temperature, such as protein-nucleic acid interactions and the oxidative state of the cell might contribute to the observed thermo-plasticity. That noted, we uncovered a sizeable excess of incompletely excised IESs that are epigenetically controlled **(Figure 3, Figure 4)**. We therefore consider it most likely that the somatic variability introduced at sub-optimal temperatures depends significantly on the environmentally induced modulation or rewiring of the epigenetic machinery regulating IES excision.

Can the observed increment in somatic genetic variation have some biological relevance? The negative correlation between IRS and fraction of exonic IESs **(Figure 5)** indicates that natural purifying selection opposes the retention of exonic IESs. This relationship is of utmost importance: it makes it very likely that somatic variability has phenotypic consequences. This interpretation is consistent with published results concerning the quality of cis-acting IES recognition/excision signals—IESs with higher quality signals lie preferentially within, rather than outside of, genes **(Ferro et al. 2015)**. It is further strengthened by the observation that non-trivial IES retention events are mainly located within weakly expressed genes **(Figure 5)**. Taken together, our findings demonstrate that part of the somatic variability in *P. tetraurelia* does not represent mere biological noise, but rather, biologically relevant selectable variation.

Are alternative somatic DNA splicing variants heritable? Our experimental setting allowed us to confidently capture trans-somatic inheritance in action. By leveraging the parallel sequencing of independently rearranged somatic genomes, we found that hundreds of somatic IESs, Dcl2/3-and Dcl5-controlled IESs, are very likely passed down to the sexual offspring after autogamy **(Figure 4)**. Additionally, the nuclear prevalence of these somatic IESs increases in the sexual progeny upon exposure to suboptimal temperatures **(Figure 3B)**. Thus, alternative DNA splicing variants may be heritable. Future studies evaluating the stability of trans-generational epigenetic inheritance of IESs across subsequent generations are required to determine e.g. whether mildly deleterious or potentially beneficial IES insertions can actually spread at the population level.

Is alternative somatic DNA splicing a source of functional innovation? To gain insight into this question, we first evaluated the impact of inefficient IES excision on gene sequences. Our observations suggest that increased IES retention following environmental perturbation result, in most cases, in the reduction of transcript availability, as inferred by the introduction of PTCs in the ORFs **(Figure 6)**. In a small fraction of observed cases, IES insertion might additionally produce diversified protein sequences **(Figure 6)**. Next, we performed an in-depth analysis of the genes hit by IES retention in response to PDE’s thermo-plasticity. We found that at least one IES-rich gene family, TPR proteins, is particularly prone/susceptible to inefficient IES excision **(Figure 7)**. Although the function of these proteins is currently unknown, their domain signatures suggest that they are involved in protein-protein interactions. Considering that TPR protein-coding genes are IES rich and yet successfully freed from IESs at 25°C, it might simply be that the excision machinery performs particularly poorly in IES rich regions at sub-optimal temperatures. Alternatively, IES excision in TPR protein-coding genes may be actively modulated in sub-optimal environments as a mechanism of gene expression control and/or to facilitate protein diversification. This alternative hypothesis is consistent with the high enrichment of epigenetically controlled IESs, Dcl5-IESs in particular, in TPR protein-coding genes (not shown). Finally, we explored the possible impact of IES retention on cellular functions and found that IES-containing genes are significantly more likely to be involved in processes such as signal transduction, cellular protein modification, and transport of ions across membranes **(Table 1)**. These results open the possibility that, much like the eukaryotic cellular process of alternative RNA splicing **(Lewis et al. 2003; Wong et al. 2013; Braunschweig et al. 2014; Marquez et al. 2015; Singh et al. 2017; Smith et al. 2018a)**, regulated IES retention in *Paramecium* might fine tune gene expression and/or generate alternative DNA splicing isoforms which may ultimately facilitate adaptation to environmental changes.

Our study offers fresh insights with regard to two current models—cryptic vs *de novo* variation-based—for explaining how selected traits become genetically encoded **(Kasinathan et al. 2017)**. Because *Paramecium* IESs are thought to be transposon-derived sequences **(Klobutcher and Herrick 1995)**, somatic IESs may be viewed both as *de novo* induced (somatic) insertions and unmasked cryptic (germline) variation at the same time. This ambiguity blurs the conventional distinction between cryptic and *de novo* induced genetic variation, rendering a discussion about the plausibility or generalizability of both models difficult. Instead, this ambiguity, together with intriguing parallels between our observations in *Paramecium* and previous findings in *Drosophila* concerning the non-random occurrence of transposon insertions in response to external stresses **(Jollos 1934; Fanti et al. 2017)**, enable cryptic and *de novo* induced variation-based mechanisms to be integrated. This exercise provides a comprehensive, potentially powerful framework for interpreting and predicting molecular dynamics associated with developmental plasticity across eukaryotes.

The framework proposed here is rooted in the idea that a number of cryptic genetic variants may be but classical genetic variants that while functional in an unstressed state, yield gene products with altered properties (e.g. down-regulated expression) in stressful environments. Under this view, a cryptic genetic variant may give rise to normally hidden phenotypes *when* it is altered in response to a stress. Such alterations, e.g., the result of stress-induced transposon insertions **(Fanti et al. 2017)** or IES retentions (this study), are at least partly nonrandom and might therefore reflect a preexisting adaptive developmental program in response to environmental adversities. This framework can account for why cryptic genetic variation may be preserved over evolutionary time. It predicts that the number of cryptic phenotypes in a population might differ based on the individuals’ levels of genetic diversity or stress susceptibility. It also incorporates observations that indicate that stress-induced genetic changes are largely epigenetically controlled, may be trans-somatically inherited, and may reach fixation epigenetically (*e.g.*, **(Sollars et al. 2003)**). Much of the stress-related genomic instability is likely to result from the disruption of small Piwi-interacting RNAs, which show several similarities between *Paramecium* and metazoans **(Bouhouche et al. 2011; Chalker and Yao 2011)**, and whose biogenesis is partially regulated by HSP90 **(Specchia et al. 2010; Gangaraju et al. 2011; Ichiyanagi et al. 2014)**.

In conclusion, we demonstrate that sub-optimal temperatures can modulate the efficiency of Programmed DNA Elimination in the ciliate *P. tetraurelia,* boosting the generation of epigenetically controlled, heritable, and functionally confined somatic DNA variability during germline-soma differentiation. The uncovered environmental sensitivity of Programmed DNA Elimination—a developmental process that unfolds in a broad range of unicellular and multicellular eukaryotes—is expected to elicit phenotypic plasticity in genetically identical organisms/cells. It also generates selectable variation, suggesting that Programmed DNA Elimination can operate as a molecular wrench that fine tunes organismal response to changing environmental conditions. Finally, our work reveals important similarities between *Paramecium* and multicellular organisms and, via these similarities, permits the elaboration of an adaptation model that is uniquely able to combine the previously argued adaptive roles of cryptic and *de novo* induced variation. Under this model, environmental cues affect the maternal environment, causing the epigenetic machinery that controls germline-soma differentiation to activate partially nonrandom plastic responses in the offspring generation. The resulting structural genomic changes are, at least partially, trans-somatically inherited and might facilitate adaptation to the triggering stress.

## Methods

### Experimental Design

To evaluate the effect of the growth temperature on PDE, fully homozygous isogenic *Paramecium* cells were cultured in daily re-isolation and passed through autogamy (self-fertilization) at three different temperatures. After a first round of autogamy at 25°C to establish a parental line, one post-autogamous parental cell was isolated and allowed to divide in fresh medium **(Figure 8**, leftmost edge). Three of the resulting isogenic cells in turn were used to start three sub-lines that were cultured in daily re-isolation and passed through a second round of autogamy at 18°C, 25°C, or 32°C to establish filial lines. Single post-autogamous F1 cells isolated for each of the sublines as well as the remaining isogenic parental cells were expanded to mass culture for somatic DNA extraction (see below).

**Figure 8.**
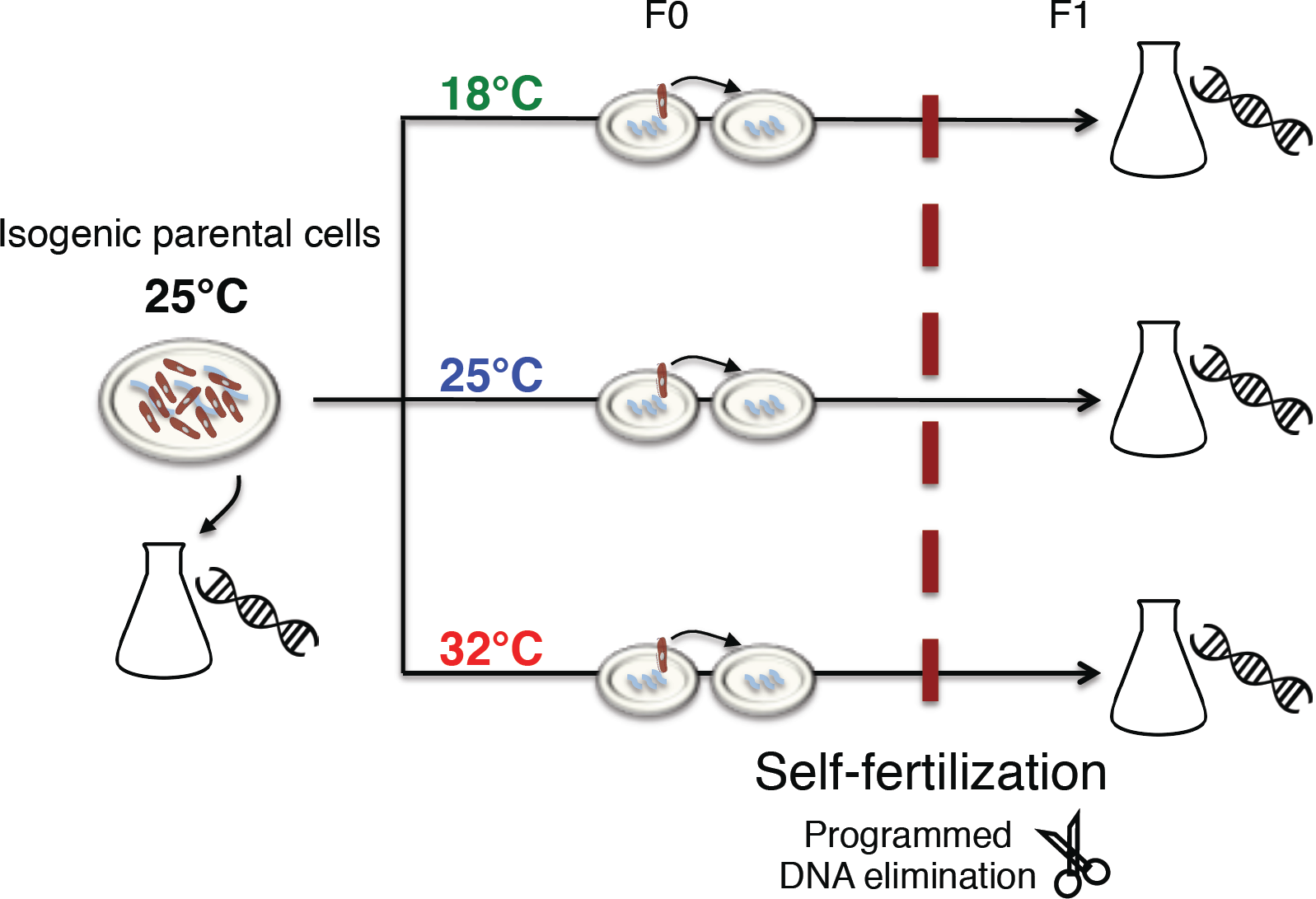
Overview of the experimental setup to characterize the impact that temperature has on the performance of Programmed DNA elimination in the ciliate *Paramecium tetraurelia* (see *Experimental design).*

### Paramecium Strains and Culture conditions

*Paramecium tetraurelia* strain d12 was used in the experiment. Cells were grown in Cerophyl Medium (CM) inoculated with *Enterobacter aerogenes* Hormaeche and Edwards (ATCC^®^ 35028TM). Stocks were fed with bacterized CM every two weeks and kept at 14°C. In preparation for the study, cells kept in stock were washed ten times in Volvic water to ensure monoxenic growth conditions during propagation. Cells were grown in depression slides under daily re-isolation regimen in 200μl of bacterized CM. Autogamy was induced by letting ∼25 division-old cells starve naturally for three days with no addition of bacterized medium. Occurrence of autogamy was confirmed by screening ≥50 Acetocarmine-stained cells for macronuclear fragmentation under a light microscope. A single ex-autogamous cell was isolated from each line, allowed to divide once, and single caryonidal progenitors brought to mass culture for somatic DNA isolation. Macronuclear fragmentation of sister caryonides was confirmed by Acetocarmine staining.

### Macronuclear DNA Isolation and Whole Genome Sequencing

The somatic nuclei of both parental (25°C_F0_) and filial lines (25°C_F1_ 32°C _F1_ and 18°C _F1_) were isolated and the macronuclear DNA (MAC) subjected to whole genome sequencing. MACs were isolated after >10 vegetative divisions post autogamy to prevent carryover of maternal MAC fragments at the time of isolation. Cells were re-suspended in Volvic water for 2 hours and allowed to digest their food vacuoles prior to MAC isolation to reduce bacterial load. MAC isolation was performed according to the protocol described in **(Arnaiz et al. 2012)**. Purified genomic DNA was subjected to ultra-deep, pair-end Illumina sequencing (∼90-100x coverage, on average, 150nt-long reads) on a HiSeq 4000 system.

### Data preprocessing

To increase the accuracy of DNA-seq data analysis raw reads were subjected to an initial step of quality control (QC) using FastQC (**http://www.bioinformatics.babraham.ac.uk/projects/fastqc/**) and pre-processed using the Joint Genome Institute (JGI) suite BBTools 37.25 **(Bushnell et al. 2017)**. Library adapters were removed from the 3’ end of the reads with BBduk using the included reference adapter file and ensuring the same length for both reads of a pair after trimming. For the k-mer based adapter detection, a long and short k-mer size of 23 and 11, respectively, were used and a single mismatch allowed. In addition, overlap-based adapter detection was enabled. Finally, short insert sizes in part of the library were leveraged for an overlap-based error correction with BBMerge, while keeping left and right reads separated.

### Calculation of IES retention scores

Quality improved, trimmed reads were mapped to the *P. tetraurelia* strain 51 reference somatic genome available via ParameciumDB **(Arnaiz and Sperling 2011)** and to a pseudo-germline genome containing all the known 44,928 IESs previously identified by Piggy MAC (PGM) knock down **(Arnaiz et al. 2012)** that was created with the Insert module of ParTIES 1.00 **(Denby Wilkes et al. 2016)**. Read mappings were performed with Bowtie 2.3.2 **(Langmead et al. 2009)** using the local alignment function for paired end reads in very sensitive mode (--very-sensitive-local) and the resulting SAM files manipulated with SAMtools 1.4.1 **(Li et al. 2009)** for downstream processing. IES retention scores (IRS) were calculated with the MIRET module of ParTIES using the IES score method.

### Genome-wide analysis of IES retention

Genome-wide analysis of IES retention was performed via statistical comparison of F0/F1 IRSs as implemented in the R script accompanying ParTIES, with the following modifications. The upper and lower bound of the 75% Confidence Interval (CI) constructed on the F0 retention score was taken as a reference retention score for binomial testing of upward or downward transitions, respectively. Lowly supported IESs i.e. IESs with a total support < 20 reads (IES+ + IES^-^) were excluded from the study. The P-values were corrected for multiple testing with the Benjamini-Hochberg method and a cutoff of 0.05 used to designate IESs with significantly different retention levels in F1 compared to F0 samples.

### Genome wide analysis of cryptic IES excisions

To estimate the rate of cryptic IES excision in response to temperature changes during nuclear differentiation TA-bound somatic deletions were characterized for the parental F0 genome and the three F1 genomes rearranged at 18°C, 25°C and 32°C using the MILORD module implemented in ParTIES. Reads mapped on the reference macronuclear genome assembly of *P. tetraurelia* strain 51 were provided as input. Low coverage cryptic IESs with total read counts (support_ref + support_variant) < 20 were excluded from the analysis. Deletion scores (DS) were calculated as the fraction of reads supporting cryptic IES excision over all reads spanning the somatic region, i.e. support_var/(support_var + support_ref). Cryptic IESs alternatively excised across samples were collected by filtering deletion scores with standard deviation > 0.3. Manual inspection with IGV **(Robinson et al. 2011)** was used to confirm putative temperature-sensitive excisions that were tagged as ‘Unstable’. Conversely, cryptic IESs consistently excised (DS > 0.5 in all samples) were tagged as ‘Stable’. The catalog of cryptic IESs is provided in **Table S1**.

### Downstream Data Analyses

We compiled a table **(Table S2**, provided as Supplementary Material) that reports the obtained IRSs of all samples and a plethora of additional IES-related information calculated *ad hoc* for this study or collected from previous studies and/or processed from various external sources, *e.g.,* the *Paramecium tetraurelia* strain 51 genome annotation v2 **(Arnaiz et al. 2017)**. All external data are publicly available at ParameciumDB **(Arnaiz and Sperling 2011)**. The following downstream analyses were conducted using in-house Python (https://www.python.org/) and R (http://www.R-project.org) scripts.

### IES-Retention Profiles Simulation

We simulated IES retention as a stochastic process. Four random samples were drawn without replacement from the full reference set of IESs (PGM-IESs). Each drawn sample contained as many elements as there were IESs with a non-zero retention score detected for each of the four DNA samples. This simulation was repeated 100 times and the maximum number of elements shared by all four drawings (theoretical 4-way overlap assuming random IES retention) used as an expectation to compare against the experimental data.

### Impact of Incomplete IES Excision on Genes

For IESs located in the coding region of protein-coding genes we checked whether their retention promotes the induction of a premature translation termination codon (PTC). We calculated the IES’s position with respect to the translation start codon, inserting the IES at this location and scanning this artificial CDS+IES-construct for an in-frame TGA (the only stop codon in *Paramecium*) upstream of the annotated one. In case a PTC was detected, we marked the distance of the PTC to the CDS start and, in case of non-3n IESs (IESs with a size that is not a multiple of 3), we marked whether the PTC occurred inside the IES body or downstream.

### Gene families and GO-term enrichment analyses

For each investigated temperature we tested whether specific gene families were over or under-represented among the genes affected by significant IES retention. We devised a sampling procedure that accounts for non-homogeneous IES densities across *P. tetraurelia*’s genes. Briefly, 10000 IES sets with size equal to the number of significantly retained IESs in the experimental samples were randomly drawn from the PGM-IESs set. A non-redundant collection of genes hit by simulated IES retention was extracted for each draw and the proportion of proteins carrying a specific functional domain (e.g. TPR, Tetratrico Peptide Repeat-containing genes, PKD, protein kinase-like domain) was used to build a null distribution. The mean of this null distribution was then taken as the success probability for a two-sided binomial test with a significance cutoff of *P* < 0.01. Raw P-values were adjusted via the Benjamini-Hochberg procedure.

The functional enrichment analysis of IES-containing genes was performed using the *statistical overrepresentation test* of the *Panther gene list analysis* tool **(Mi et al. 2013; Mi et al. 2017)** and cross-validated with the *topGO* package **(Alexa and Rahnenfuhrer 2016)**. The mRNA IDs (e.g. GSPATT00000013001) of lES-containing genes (macronuclear gene models v1) were used as supported gene identifiers for the *Panther gene list analysis.* In all cases, IES-containing genes were tested against the full set of macronuclear (coding) gene models of *P. tetraurelia.* The Weigh01 algorithm and F statistic (Fisher’s exact test) were used for testing the GO-terms overrepresentation with the *topGO* package. Raw P-values provided by *topGO* were adjusted with the *P.adjust* function (method = “hochberg”) implemented in the R package *stats* (version 3.4.0). A critical value of 0.05 was adopted as significance threshold in all tests.

### Data availability

All the DNA sequence reads obtained for control and temperature exposed samples have been deposited at the European Nucleotide Archive: PRJEB28697.

## Acknowledgements

We thank Gennady Churakov, Franz Goller, and Hans-Dieter Görtz for their valuable comments on a draft of the manuscript. Kathrin Brüggemann is gratefully acknowledged for her technical assistance. This work was supported by a Deutsche Forschungsgemeinschaft (DFG) research grant to FC [CA1416/ 1-1] and carried out within the DFG Research Training Group 2220 ‘Evolutionary Processes in Adaptation and Disease’ at the University of Münster.

## Author contributions

Valerio Vitali: *Investigation, Formal analysis, Software, Methodology, Visualization, Writing-original draft, Writing-review & editing.* Rebecca Hagen: *Data curation, Software, Writing-review & editing.* Francesco Catania: *Funding acquisition, Conceptualization, Project administration, Supervision, Visualization, Writing-original draft, Writing-review & editing.*

